# Dynamic modulation of glucose utilisation by glucocorticoid rhythms in health and disease

**DOI:** 10.1101/2020.02.27.968354

**Authors:** Eder Zavala, Carlos A. Gil-Gómez, Kyle C. A. Wedgwood, Romana Burgess, Krasimira Tsaneva-Atanasova, Marco A. Herrera-Valdez

## Abstract

A systems level coordination of physiological rhythms is essential to sustain healthy states, especially in the face of stimuli that may disrupt such rhythms. The timing of meals, medication and chronic stress can profoundly influence metabolism, which depends on the dynamic interactions between glucose, insulin and cortisol. Although the metabolic and stress endocrine axes are simultaneously disrupted in many diseases, a theoretical framework to understand how chronodisruption leads to disease is lacking. By developing a mathematical model of glucose utilisation that accounts for the antagonism between insulin and cortisol, we investigate the dynamic effects of glucose boluses under normal and disrupted cortisol rhythms, including the effects of cortisol agonists and antagonists. We also predict how cortisol rhythms modulate circadian responses to oral glucose diagnostic tests, and analyse the disruptions caused by hypercortisolism. Finally, we predict the mechanisms leading to type 2 diabetes in patients with normal and excess cortisol.

## Introduction

Glucocorticoids are vital steroid hormones that mediate the stress response, modulate glucose metabolism, and have anti-inflammatory and cognitive effects. Key to their physiological function, circulating levels of glucocorticoids such as cortisol (CORT) fluctuate with circadian and ultradian periodicity, reaching maximal values during the waking period and progressively decreasing during the day before rising again at night (*1*). Disruption of these hormone rhythms often leads to disease (*2*), with chronic CORT excess (hypercortisolism) commonly associated with the impairment of glucose metabolism and the development of secondary type 2 diabetes (*3, 4*). This is the case in both endogenous and medication-induced hypercortisolism (Cushing’s syndrome), with up to 70% of patients developing impaired glycaemic control (*4*). In addition, chronic conditions such as coronary artery disease, fibromyalgia, rheumatoid arthritis, and mood disorders such as Major Depressive Disorder (MDD) and Post Traumatic Stress Disorder (PTSD) have also been associated with abnormally high CORT levels and metabolic disruptions (*5-13*). Interestingly, while diabetes in people with normal CORT levels is characterised by increased fasting glucose, a static metric by all means, secondary diabetes in hypercortisolemic patients can be better characterised by impaired glucose tolerance, a medical test that is intrinsically dynamic (*14, 15*). However, a theoretical framework to help understand the complex regulation that fluctuating stress hormones exert on glucose homeostasis is lacking.

Although most mathematical models of metabolic disorders focus on understanding the pathogenesis of type 2 diabetes, these models typically ignore the effects of CORT dynamics. Recent work has recognised the importance of understanding the impact of chronodisruption on glucose metabolism (*16*), and several models (*17-19*) have re-examined physiological and clinical data from a dynamical perspective to shed light on the mechanisms leading to diabetes, with uncompensated insulin resistance at their core (*20, 21*). The work of Ha & Sherman, for example, proposes mathematical models accounting for key mechanisms mediating beta-cell responses to hyperglycaemia, namely: increased sensitivity and increased secretory response over short and intermediate timescales, and changes in beta-cell mass over longer timescales (*19, 22*). By systematically exploring the contribution of these mechanisms, each represented by a specific parameter in their model, the authors predict thresholds that mark the transition from pre-diabetes to diabetes, supporting some of the hypotheses already advanced by (*21*) and in agreement with clinical observations. Although the pathogenesis of secondary diabetes under hypercortisolism is still unclear, some key driving mechanisms have been identified such as the stimulation of gluconeogenesis and development of insulin resistance, together with the impairment of pancreatic insulin secretion (*23, 24*). Nevertheless, the effect of chronic CORT excess on these regulatory mechanisms has remained unexplored.

In this work, we present a mathematical model of glucose homeostasis that accounts for CORT dynamics and predicts the dysregulation elicited by hypercortisolism in the onset of type 2 diabetes. By considering the mechanisms mediating insulin and CORT antagonism, we explore the effects of circadian and ultradian CORT rhythms on the regulation of blood glucose. We first validate our model against data from oral glucose tolerance tests (OGTTs) in healthy subjects, and predict the dynamic effects of OGTTs following sub-chronic treatment by glucocorticoid agonists and antagonists of clinical relevance. Secondly, we use our model to investigate the effects of ultradian and circadian rhythms in the interpretation of OGTTs used for diagnosis, and how hypercortisolism affects circadian variability of fasting glucose and insulin, as well as their response following OGTTs. Finally, we contrast our model with longitudinal data from OGTTs in diabetics to 1) evaluate the relative contribution of disrupted regulatory mechanisms in the progression of diabetes in people with normal CORT levels, and 2) to predict the glucose and insulin responses following OGTTs observed in the progression of diabetes in patients with hypercortisolism. We discuss our results in the light of circadian timing approaches to interpret clinical data (chronodiagnosis), and in the design of chronotherapies for the metabolic complications associated to stress hormones.

## Materials and Methods

### Mathematical modelling

Current models of the pathogenesis of diabetes typically place the feedback interactions between glucose and insulin at the core of glycaemic control (*17, 25*). While these models assume that blood glucose uptake into cells directly depends on circulating levels of insulin, the mechanisms by which insulin regulates transmembrane glucose transport have not been explicitly modelled. Although ignoring such mechanisms significantly simplifies the computational analysis, it also leaves out important effects such as the differential sensitivity to glucose across tissues and the antagonism between glucocorticoids and insulin. In our model (Fig. 1), we take a step toward addressing this by accounting for the effect of transmembrane glucose transporters (GLUTs) in mediating glucose uptake from blood into fat and skeletal muscle cells, a process that is amplified by the translocation of GLUTs from intracellular pools to the cell membrane following insulin stimulation (*26, 27*). GLUTs are also involved in glucose sensing in pancreatic beta cells, with the caveat that the GLUT isoforms present in these insulin-secreting cells possess a different glucose sensitivity threshold than those in fat and muscle cells (*28-31*). Furthermore, CORT regulates glucose uptake by antagonising the insulin-mediated translocation of GLUTs in fat and muscle (*32-35*), by modulating insulin secretion in pancreatic beta cells (*36-39*), and by stimulating liver gluconeogenesis and glycogenolysis (*4, 15*). Importantly, our model considers the distribution of GLUT isoforms with different glucose affinities for glucose transport across cell membranes (Supplementary Material). While other organ tissues also exhibit GLUT mediated glucose uptake, we focus on blood glucose clearance in fat and muscle given that this process is simultaneously regulated by insulin and CORT. Thus, we postulate a model that considers how surges in blood glucose (induced by food intake or OGTTs) elicit insulin secretion, which in turn increase the fraction of active GLUT transporters in fat and muscle cells, resulting in the import of glucose from blood into these cells (Fig. 1). Importantly, CORT antagonises this process *dynamically*, and we model it as an external driving force with ultradian periodicity and circadian amplitude. Similarly, we consider OGTTs as single perturbations that elicit a transient dynamic response depending on the dynamics of CORT at the time of the pulse.

**Fig. 1.**
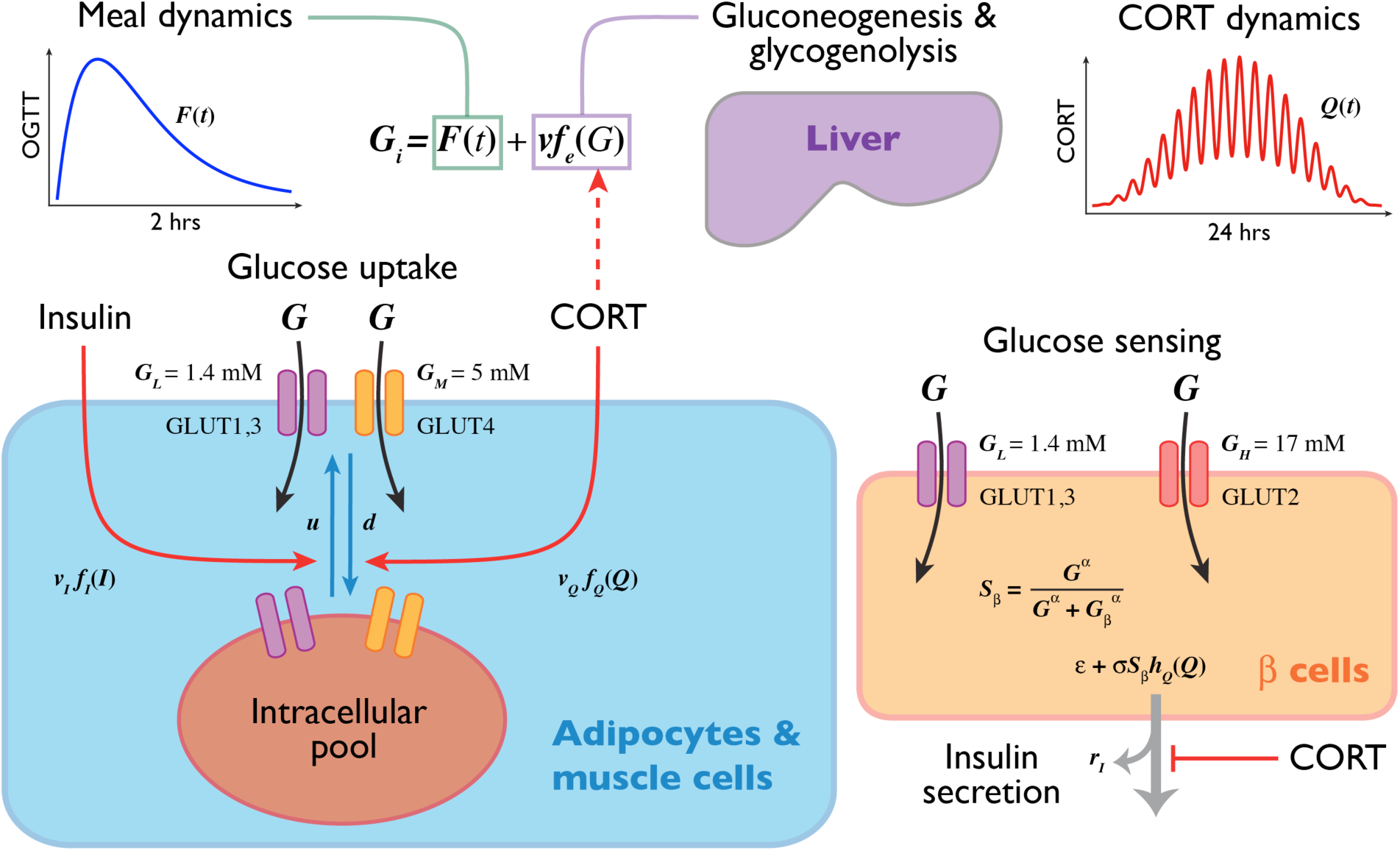
Pictorial summary of the model. Blood glucose levels are primarily controlled by food intake, liver gluconeogenesis, glycogenolysis, and utilisation by fat and skeletal muscle. Pancreatic beta cells continuously sense circulating levels of glucose to secrete insulin. Glucocorticoids modulate these processes dynamically by antagonising insulin-mediated translocation of GLUTs in fat and muscle, inhibiting pancreatic insulin secretion and, in hypercortisolism, stimulating gluconeogenesis and glycogenolysis.

Regarding insulin secretion, it is known that systemic administration of glucocorticoids induces insulin resistance, followed by a compensatory increase of beta cell mass and ultimately, insulin secretory failure (*40*). However, assessing the effects of glucocorticoids on beta cell function is challenging because of the abundance of apparently conflicting reports. These arise because glucocorticoid effects depend on the dose, potency, and duration of exposure of the steroid used, the cellular context (e.g., *in vivo* vs. *in vitro*) and the susceptibility of the cell line (*37*). We chose to model CORT as inhibiting insulin secretion in beta cells as this effect occurs following exposure to glucocorticoids within a short timescale, which is relevant to the timescale of ultradian CORT oscillations and OGTTs dynamics (*38*).

Lastly, we also consider endogenous glucose release from the liver (*24*), a process that we modelled as active primarily during fasting but diminished following glucose intake from meals or OGTTs. This process, together with pancreatic insulin secretion and insulin effects in fat and muscle cells was considered in our model as being dynamically regulated by CORT. The dynamics of CORT was modelled as ultradian oscillations (75 min period) with circadian modulated amplitude and baseline reaching a maximum CORT awakening response at 7 am (*41*) (Fig. 1 and Supplementary Material).

### Computer simulations and parameter estimation

The model equations were solved and analysed using Python 3.6 and XPPAUT (*42*). Details of the mathematical model development and parameter estimation are described in the Supplementary Material. Computer code to run *in silico* experiments is available on GitLab at https://gitlab.com/ezavala1/modelling-cortisol-dynamics-in-glucose-homeostasis.

The model parameters were estimated from the literature and recalibrated by fitting the model output to data from OGTTs in healthy humans (*43*) and to clinical data from OGTTs in diabetics (*21*). Baseline parameter values were estimated as follows: 1) the beta-cell secretory function was fitted to data from glucose-induced insulin secretion in perfused human islets (*44, 45*); 2) the kinetic rates involved in the translocation of GLUT transporters, including the effects of insulin and glucocorticoids, were estimated from (*27, 34, 46*); 3) the parameters associated to glucose uptake in peripheral cells, as well as maximum rates of production and utilisation, were estimated from (*31, 43, 44*). The remaining parameters were initially set to biologically plausible values and recalibrated during model fitting. A table of parameter values and their source is available in the Supplementary Material.

## Results

### Cortisol dynamics modulates responses to OGTTs

We first investigated the ability of our model to reproduce OGTTs in humans. To do this, we drove glucose and insulin levels in the model with CORT rhythms and simulated a pulse of glucose equivalent to an OGTT administered at 8 am. The model predicts dynamic transient responses that match continuously sampled levels of glucose and insulin in blood every 10 min during the first 30 min, and then every 30 min until 3 hrs following the glucose bolus (*43*) (Fig. 2A).

**Fig. 2.**
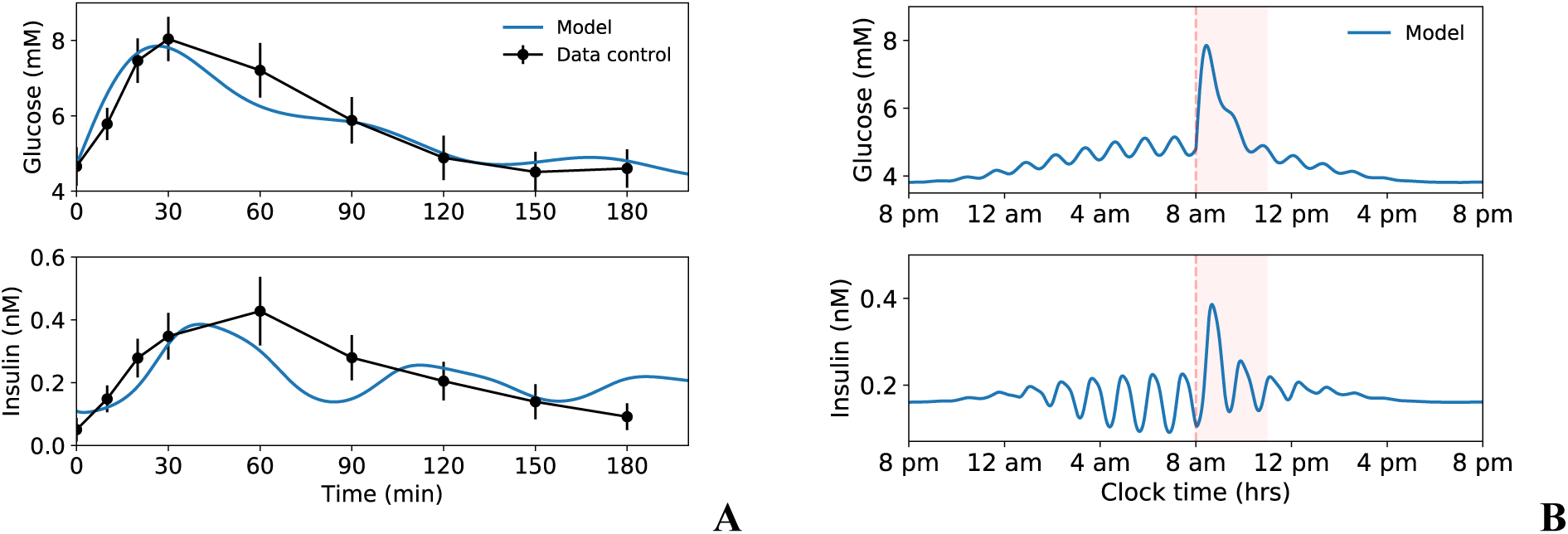
Transient dynamic responses following simulation of OGTTs. **A.** Comparison of the model dynamics against data from OGTTs in humans (*43*), predicting transient dynamic increases in glucose and insulin, as well as ultradian oscillations driven by CORT. **B.** 24 hr dynamics of glucose and insulin including a transient response following an OGTT.

Numerous studies have reported the presence of ultradian oscillations of insulin and glucose. It is no surprise that the reported frequency of these oscillations usually varies with the sampling rate, pulse detection method, and measuring conditions (e.g., *in vivo* vs. *in vitro*). In addition, work from Peter Butler’s lab (*47, 48*) and pancreatic islets measurements (*49, 50*) suggests an attenuation of the 4-20 min insulin pulsatility at the portal vein due to extraction and dilution as it progresses through the systemic circulation. Given the systems-level scope of our model, we predict slow ultradian oscillations (∼80 min period) arising from the CORT-modulated feedback loop between glucose and insulin (Fig. 2B). This result agrees with previously reported ultradian oscillations (80-150 min period) of glucose and insulin measured in healthy subjects, where the origin of these slow ultradian oscillations was first hypothesized (*51*). The model also predicts ultradian oscillations in the fraction of active GLUTs and their associated transient dynamic responses following OGTTs, although with a smaller magnitude (Supplementary Material). According to the model, these transient dynamic responses last between 2-3 hrs following the glucose bolus (Fig. 2B).

### OGTT responses are modulated by glucocorticoid agonists and antagonists

We next used the model to investigate the effects that sub-chronic administration of agonists and antagonists of glucocorticoids have on the insulin and glucose dynamic response to an OGTT. These drugs are widely used clinically for their action on the hypothalamic-pituitary-adrenal axis and the regulation of immune responses, including anti-inflammatory effects. For example, it has long been known that prolonged administration of dexamethasone, a powerful agonist of the glucocorticoid receptor, inhibits secretion of endogenous CORT (*52*). At these days-long timescales, dexamethasone increases insulin secretion (*36, 43*) and inhibits the translocation of GLUTs to the cell membrane in response to insulin (*53*). Including these regulatory effects into our mathematical model (Supplementary Material) predicts a larger transient dynamic response of glucose and insulin following OGTTs (Area Under the Curve (AUC)_G-Dex_ = 1255.85, AUC_I-Dex_ = 71.5) than without dexamethasone treatment (AUC_G_ = 1050.41, AUC_I_ = 39.26). This prediction agrees with data from OGTTs in humans following 5-day treatment with dexamethasone (*43*) (Fig. 3A). Notably, the model also predicts the loss of ultradian oscillations of glucose and insulin following the OGTT perturbation, possibly because of the suppression of endogenous CORT rhythms elicited by sub-chronic dexamethasone.

**Fig. 3.**
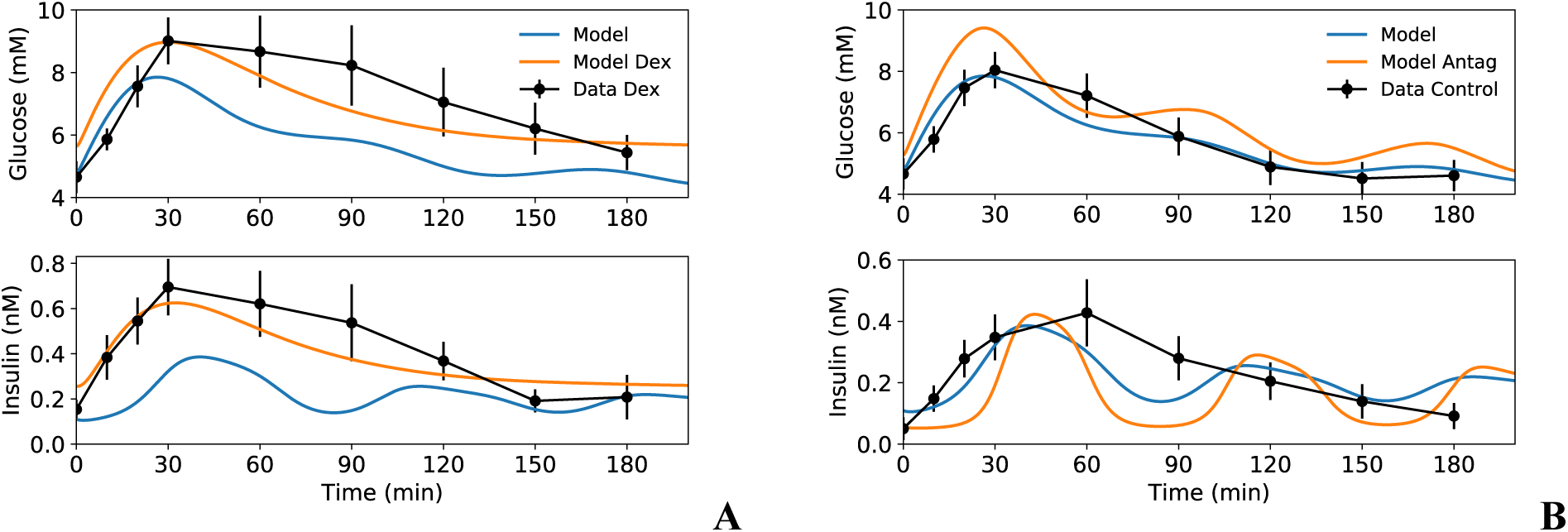
Treatment with CORT agonists and antagonists modulates glucose and insulin response to OGTTs. **A**. The model predicts an enhanced glucose and insulin non-oscillatory OGTT response following sub-chronic treatment with the CORT agonist dexamethasone (*43*). **B**. The model predicts an enhanced glucose response following an OGTT under sub-chronic treatment with the CORT antagonist Org34850 (*54*). It also predicts ultradian oscillations of glucose and insulin with higher amplitude than in the absence of the antagonist (data from (*43*)).

The model also predicts a transient dynamic response to an OGTT following sub-chronic treatment with a CORT antagonist (*54*) (Supplementary Material). Interestingly, the predicted response not only retains ultradian oscillations in glucose and insulin but the amplitude of these oscillations is higher than without treatment with the antagonist, albeit with a reduced cumulative insulin secretion (AUC_G-Ant_ = 1182.04, AUC_I-Ant_ = 29.33) (Fig. 3B). To the best of our knowledge, this model prediction has not been experimentally confirmed. However, our results suggest that a high sampling rate (roughly every 10 min) of glucose and insulin following an OGTT will be necessary to detect the oscillatory dynamics predicted by the model.

### Circadian and ultradian sensitivity to OGTTs in normal physiology and hypercortisolism

Understanding the implications of circadian timing in the interpretation of diagnosis tests is a major challenge for the accurate diagnosis and treatment of metabolic and endocrine conditions. These tests typically involve perturbing the system by administering a drug, a hormone, a toxin, or a nutrient, and measuring the resulting dynamic response over time. The OGTT is arguably the most commonly used test for diagnosing and monitoring the progression of abnormal glycaemia. Thus, we used our mathematical model to investigate the dependency of the transient dynamic response elicited by OGTTs on the circadian and ultradian phase of CORT. To do this, we simulated a single OGTT administered at different time points over a 24-hr period (∼2 min between time points) and recorded the magnitude of the glucose and insulin response measured between the baseline at the time of the OGTT and the maximum peak post-OGTT. The result is shown in Fig. 4A, where clear circadian and ultradian patterns emerge in the magnitude of the glucose and insulin responses. Consistent with clinical and experimental observations (*55, 56*), the model predicts peak responses during the inactive phase of the 24h cycle during which circadian CORT levels increase, and are minimal during the active phase, when CORT levels decrease. Of importance to interpret OGTTs, our model also predicts different glucose and insulin responses when we look at the effects of ultradian rhythmicity. While the glucose response peaks when the OGTT arrives during the falling phase of the ultradian cycle (*π*/2 to *π*), the insulin response peaks when the OGTT arrives during the early rising phase (0 to *π*/4). Interestingly, the ultradian dependency is minimal for both glucose and insulin between 12 pm and 10 pm, indicating that responses to an OGTT in the afternoon are blind to whether the glucose bolus arrives at the rising or falling phase of the ultradian rhythm.

**Fig. 4.**
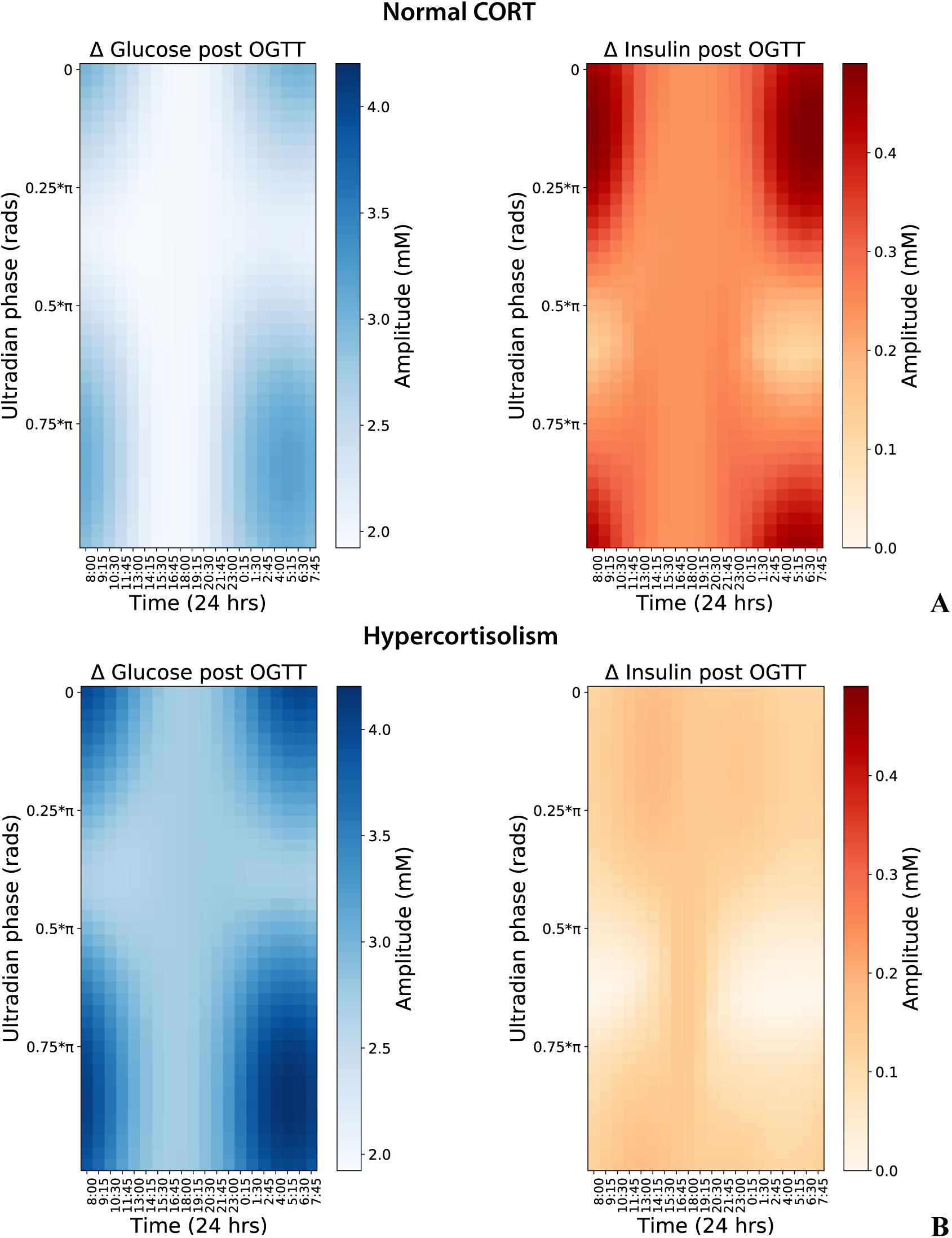
Circadian and ultradian variability of glucose and insulin responses to OGTTs. **A.** Under normal levels of CORT, glucose and insulin responses to OGTTs show circadian variability, with maximal values during the night and minimal during the day. The model also predicts ultradian variability, with glucose and insulin showing maximal responses to OGTTs arriving, respectively, during the falling and rising phase of the ultradian rhythm. **B.** Excess CORT increases the magnitude of the glucose responses to OGTT while preserving circadian and ultradian variability, but significantly inhibits insulin responses as well as their circadian and ultradian variability.

We then asked the question of whether hypercortisolism affects the circadian and ultradian responses of glucose and insulin to OGTTs. First, it is important to stress that in hypercortisolism, it is not only that CORT levels are higher than normal, but in some cases, they also display abnormal rhythmicity. This varies depending on whether the underlying condition is associated with Cushing’s syndrome, depression, PTSD, acute inflammation following major surgery, or chronic inflammatory diseases (e.g., rheumatoid arthritis) (*2, 5-13, 57-60*). Secondly, many of these conditions are accompanied by a disruption of normal glucose metabolism while preserving some degree of CORT rhythmicity. Since we want to investigate solely the effects of CORT excess rather than its combination with disrupted rhythmicity observed within each condition, we modelled hypercortisolism by approximately doubling normal physiological CORT levels while preserving normal ultradian and circadian periodicity (Supplementary Material). The results are shown in Fig. 4B, where we can see that glucose responses to OGTTs maintain the same qualitative dependency on ultradian and circadian rhythms as in the normal CORT regime, but with significantly enhanced amplitude. Conversely, insulin responses significantly decrease in amplitude and lose their circadian and ultradian variability under hypercortisolism. These effects are likely due to the direct CORT inhibition of pancreatic insulin secretion, which in turn induces exacerbated glucose responses following OGTTs.

### Hypercortisolism disrupts the mechanisms underpinning glucose-insulin homeostasis

Understanding the origin of diurnal rhythms in glucose and insulin is challenging due to the diversity of physiological processes that simultaneously modulate such dynamics. These range from the temporal distribution of feeding (*56*), upstream control via the suprachiasmatic nucleus (*61*), circadian skeletal muscle insulin sensitivity and glucose uptake (*62*), and circadian clock genes (*63, 64*). Not surprisingly, uncovering the mechanisms underlying chronodisruption pose additional challenges, mostly because it is difficult to identify unequivocally what mechanisms break down, in what order, and how effects accumulate to lead to disease. In the context of circadian modulation of glucose metabolism, it has been recognised that understanding how the disruption of such mechanisms is deleterious for health is key to develop appropriate therapeutic interventions (*65*). Thus, we used our mathematical model to investigate the contribution to circadian variability of physiological processes within the glucose-insulin systems-level feedback loop.

As in the previous section, we analysed glucose and insulin responses following OGTTs. In this case, we assumed all OGTTs were given at the minima of the ultradian rhythm (phase *φ*= 0 = *π* in Fig. 4A), and recorded both the baseline and the magnitude of the response following an OGTT. We did this by simulating OGTTs occurring at different times of the day across the 24h cycle, under normal CORT levels and hypercortisolism, and testing the effect of changing different parameter values representing physiological mechanisms known to be disrupted in the development of type 2 diabetes (Fig. 5). In a first instance, we considered the simplest scenario where no such mechanism was disrupted and hence no parameters other than those associated with the effects of CORT excess were varied (Fig. 5A). We quantify the effects of hypercortisolism and of changing parameter values by calculating the percentage change in the AUC of basal and post-OGTT glucose and insulin throughout the day with respect to nominal parameter values. Consistent with previous simulations, the model predicts circadian variability of basal glucose and its response to OGTTs, which is not disrupted by hypercortisolism but only shifted towards higher values (ΔAUC_G-basal_ ≈ 27%, ΔAUC_G-OGTT_ ≈ 35%). In contrast, insulin responses reduce in magnitude (ΔAUC_I-OGTT_ ≈ −57%) and lose their circadian variability under hypercortisolism, while their baseline levels maintain their circadian variability but decrease in magnitude (ΔAUC_I-basal_ ≈ −48%) (Fig. 5A).

**Fig. 5.**
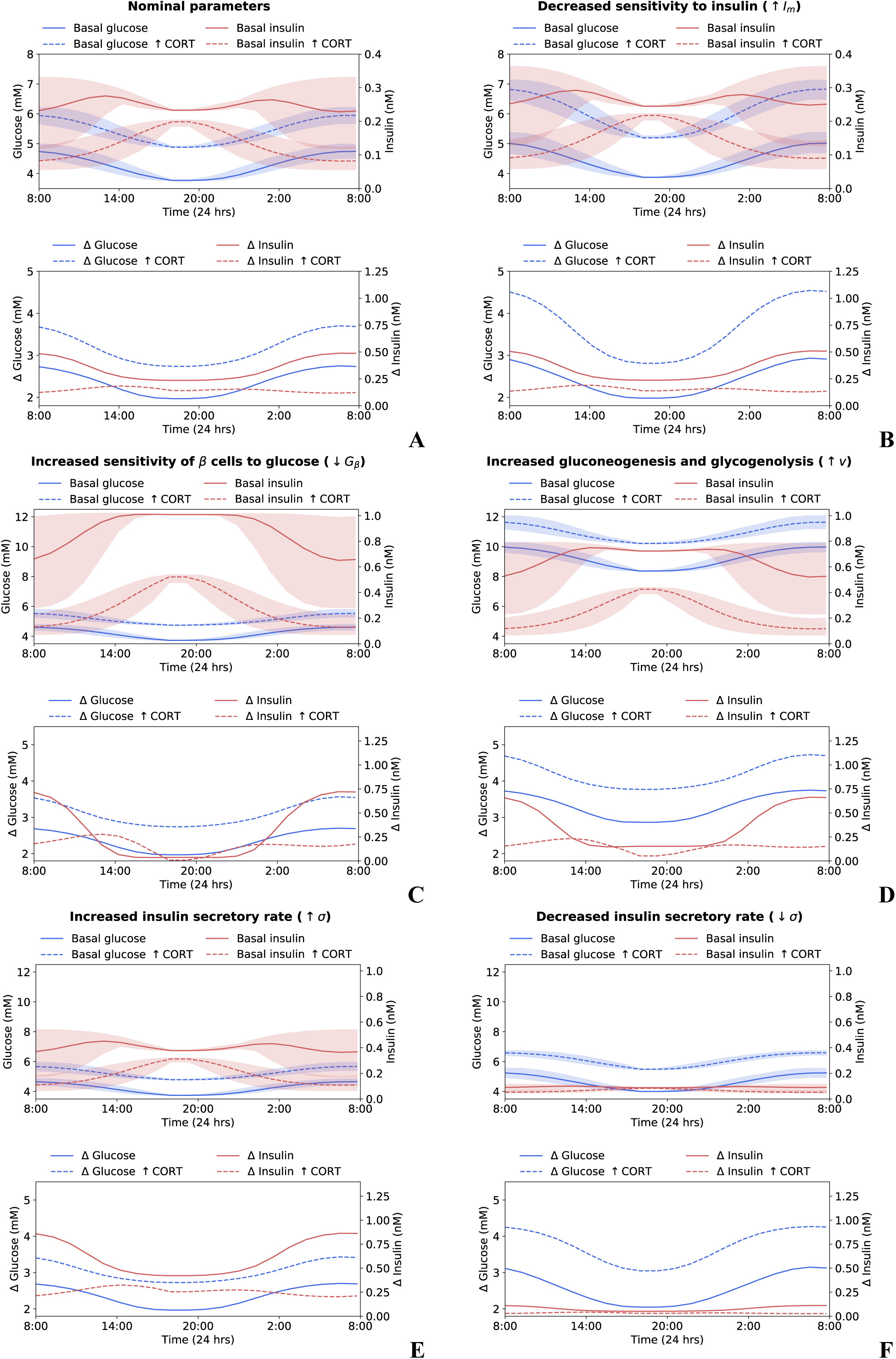
Circadian and ultradian variability of glucose and insulin at baseline and post-OGTT responses. The variability was assessed by simulating an OGTT at different times of the day but administered at the nadir of the ultradian rhythm. Shaded areas indicate the range of the ultradian rhythm. Both normal CORT and hypercortisolism cases were investigated for six different scenarios: **A.** All mechanisms underlying glucose homeostasis are normal. **B.** Decreased muscle and fat cell sensitivity to insulin. **C.** Increased sensitivity of beta cell secretory response to glucose. **D.** Increased gluconeogenesis and glycogenolysis. **E.** Increased insulin secretion in pre-diabetes. **F.** Decreased insulin secretion in diabetes.

In the development of secondary type 2 diabetes, there are several mechanisms that could be potentially disrupted by CORT excess. Thus, we used the model to test the effects of hypercortisolism on the circadian and ultradian variability of glucose and insulin fasting levels and their response to OGTTs when each of these mechanisms were disrupted by CORT. To do this, we changed the value of model parameters accounting for these mechanisms independently, and tested both scenarios with normal CORT and hypercortisolism. The results are shown in Figs. 5B-F where, respectively, we lower the muscle and fat cell sensitivity to insulin (Fig. 5B), increase the sensitivity of beta cell secretory response to glucose (Fig. 5C), increase the rate of gluconeogenesis and glycogenolysis (Fig. 5D), increase the insulin secretory rate associated with pre-diabetes (Fig. 5E), and decrease the insulin secretory rate associated with diabetes (Fig. 5F).

We found that lowering the muscle and fat cell sensitivity to insulin (*I*_*m*_ = 0.08 to 0.32 nM) (Eq. 10 in the Supplementary Material) slightly increased the basal levels of glucose and insulin (ΔAUC_G-basal_ ≈ 4%, ΔAUC_I-basal_ ≈ 6%), and that hypercortisolism slightly amplified this effect (ΔAUC_G-basal_ ≈ 11%, ΔAUC_I-basal_ ≈ 11%) (Fig. 5B). Similarly, glucose and insulin responses to OGTTs remain largely unchanged (ΔAUC_G-OGTT_ ≈ 3%, ΔAUC_I-OGTT_ ≈ 2%), with the only notable effect induced by hypercortisolism was to increase the amplitude of the circadian variability in glucose responses (ΔAUC_G-OGTT_ ≈ 15%, ΔAUC_I-OGTT_ ≈ 5%).

The effect of modulating the beta cell sensitivity to glucose has previously been investigated through mathematical modelling (*19*). Here we take forward the analysis to include circadian variability on basal glucose, insulin, and their post-OGTT responses. We find that increasing the sensitivity of beta cell secretory response to glucose (*G*_*β*_ = 6.48 to 2.59 mM) (Eq. 9 in the Supplementary Material) dramatically increases basal insulin (ΔAUC_I-basal_ ≈ 256%) and the circadian variability of its ultradian rhythm, while the effects on basal glucose were barely noticeable (ΔAUC_G-basal_ ≈ −2%) (Fig. 5C). Regarding the OGTT responses, it is also insulin that shows increased circadian variability, with larger responses between 4 am and 12 pm, and smaller ones between 2 pm and 12 am, resulting in a reduced cumulative dose during the day (ΔAUC_I-OGTT_ ≈ −13%). Interestingly, under increased beta cell sensitivity to glucose, hypercortisolism decreased basal glucose levels only slightly (ΔAUC_G-basal_ ≈ −6%), while notably increased basal insulin (ΔAUC_I-basal_ ≈ 119%). CORT excess also increased the circadian variability of insulin responses (ΔAUC_I-OGTT_ ≈ 7%), while the amplitude of glucose responses remained largely unchanged (ΔAUC_G-OGTT_ ≈ −2%).

Next, we tested the effect of increasing the endogenous glucose production rate corresponding to gluconeogenesis and glycogenolysis (*v* = 18.69 to 37.38 mM min^-1^) (Eq. 4 in the Supplementary Material). This parameter had by far the most noteworthy effect on increasing glucose baseline levels and its responses to OGTTs (ΔAUC_G-basal_ ≈ 116%, ΔAUC_G-OGTT_ ≈ 41%) (Fig. 5D). Hypercortisolism amplified these effects (ΔAUC_G-basal_ ≈ 101%, ΔAUC_G-OGTT_ ≈ 32%) while preserving the circadian variability. Increasing the endogenous glucose production also increased basal insulin (ΔAUC_I-basal_ ≈ 169%), while the magnitude of its response to OGTTs remained unchanged (ΔAUC_I-OGTT_ ≈ 1%) albeit with increased circadian variability (Fig. 5D). Hypercortisolism produced only a small change in the circadian variability of post-OGTT insulin responses, with a cumulative dose largely unchanged (ΔAUC_I-OGTT_ ≈ 2%).

Finally, we tested the effect of an increased insulin secretion rate observed in the early stages of glucose metabolism disruption, and a decreased insulin secretion rate observed subsequently in diabetes (Fig. 5E-F). As expected, an increased insulin secretion rate (*σ* = 0.48 to 1.34) (Eq. 2 in the Supplementary Material) increased baseline insulin levels and its responses to OGTTs (ΔAUC_I-basal_ ≈ 62%, ΔAUC_I-OGTT_ ≈ 78%), while baseline and post-OGTT glucose levels did not change (ΔAUC_G-basal_ ≈ −1%, ΔAUC_G-OGTT_ ≈ −1%) (Fig. 5E). Under hypercortisolism, baseline glucose levels, their post-OGTT responses and their circadian variability were largely unchanged (ΔAUC_G-basal_ ≈ −4%, ΔAUC_G-OGTT_ ≈ −4%), while insulin baseline levels and its responses increased (ΔAUC_I-basal_ ≈ 47%, ΔAUC_I-OGTT_ ≈ 76%). In contrast, decreasing the insulin secretion rate (*σ* = 0.48 to 0.19) notably reduced both insulin basal levels, its post-OGTT responses, and erased their circadian variability (ΔAUC_I-basal_ ≈ −63%, ΔAUC_I-OGTT_ ≈ −78%) (Fig. 5F). Furthermore, glucose basal levels and post-OGTT responses remained within normal ranges and preserved their circadian variability (ΔAUC_G-basal_ ≈ 8%, ΔAUC_G-OGTT_ ≈ 8%). Simulating the hypercortisolism scenario also reduced insulin levels and its responses (ΔAUC_I-basal_ ≈ −50%, ΔAUC_I-OGTT_ ≈ −78%), while it increased both glucose basal levels and post-OGTT responses (ΔAUC_G-basal_ ≈ 13%, ΔAUC_G-OGTT_ ≈ 17%).

### Predicting excess CORT induced disruption of glucose and insulin dynamics

One of the main challenges in understanding the mechanisms underpinning glucose homeostasis consists of untangling their synergistic action. This is particularly important considering that both their sequential or simultaneous disruption can lead to manifestations in abnormal glucose and insulin dynamics, often resulting in the development of type 2 diabetes. As shown in the previous section, our mathematical model predicts how these mechanisms influence circadian variability of glucose and insulin dynamics, and how excess CORT induces chronodisruption. However, it is also important to assess the contribution of each mechanism to the disruption of insulin and glucose dynamics as disease develops. Thus, we now use the mathematical model to explore the combined contribution of disrupting mechanisms regulating insulin and glucose utilisation. To do this, we start by considering normal CORT levels and validate our model predictions against data from diabetic patients at different stages of disease progression, from the early development of glucose intolerance and hyperinsulinemia to hyperglycaemia and hypoinsulinaemia (*21*). These clinical observations are represented through dynamic changes in glucose and insulin responses to OGTTs vs. fasting glucose levels, and highlight the observation of increased insulin secretion in prediabetics compared to healthy subjects known as Starling’s law of the pancreas (*19*). In the model, we chose combinations of model parameters as described in Fig. 5 and ran computer simulations to determine the parameter values that predict insulin and glucose responses to OGTTs for increasing baseline glucose levels. By successively mapping and fitting these values to Starling’s curve, we identified three mechanisms with the largest contribution to the onset and progression of type 2 diabetes. These mechanisms are represented by model parameters *σ* (insulin secretion rate), *v* (endogenous glucose production rate due to gluconeogenesis and glycogenolysis), and *I*_*m*_ (muscle and fat cell sensitivity to insulin). Of importance, we only allowed the insulin secretion rate *σ* to increase for relatively low values of fasting glucose and to decrease for higher fasting glucose levels (threshold set at ∼6.3 mM). These results are summarised in Fig. 6 (green curves), where we show that insulin responses increase with fasting glucose levels before reaching the Starling’s curve peak, after which insulin responses decrease while the subject develops hyperglycaemia.

**Fig. 6.**
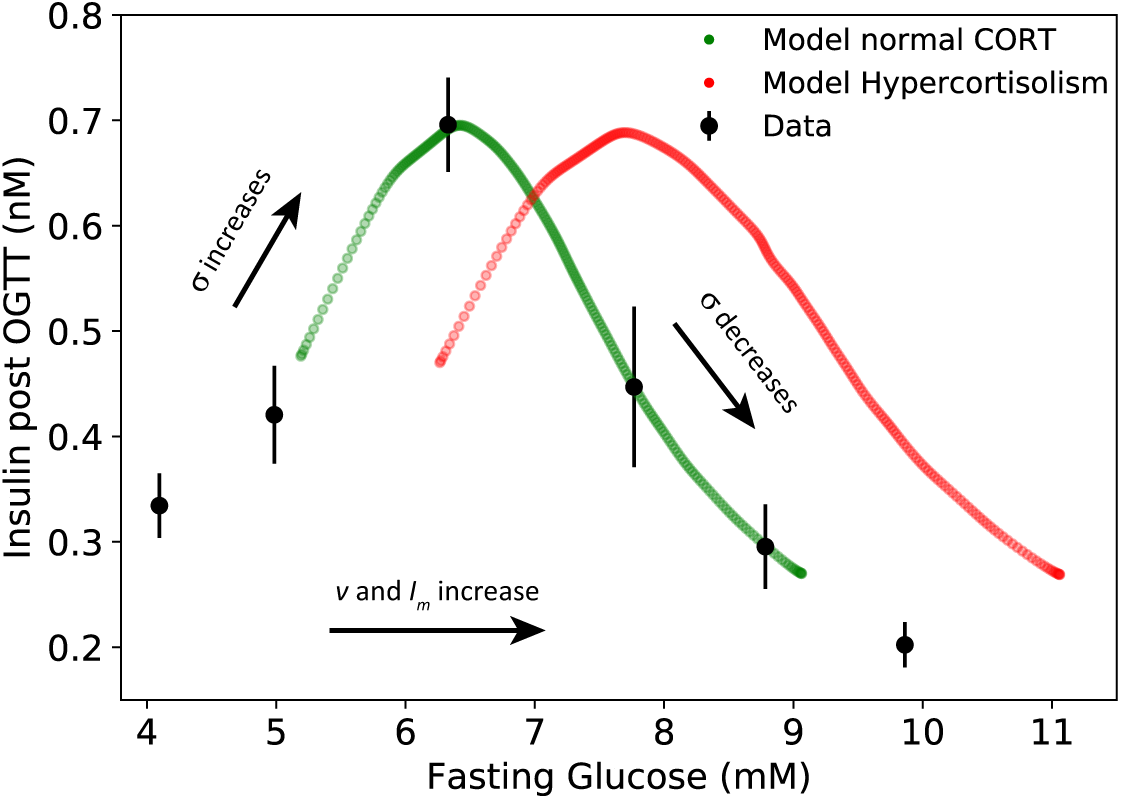
Predicting paths to diabetes in normal CORT vs. excess CORT patients. Disruptions of insulin responses are described by Starling’s law of the pancreas, characterised by an increase in insulin secretion for relatively low fasting glucose (pre-diabetes), followed by a decrease in insulin secretion for high fasting glucose (diabetes) (*21*). The model predicts how the disruption of insulin responses to OGTTs is driven by simultaneous changes in the insulin secretion rate *σ*, endogenous glucose production rate *v*, and muscle and fat cell sensitivity to insulin *I*_*m*_. Our model also shows how excess CORT shifts this path to higher fasting glucose levels.

Next, we used the mathematical model to predict the disruption of glucose and insulin dynamics in the onset and progression of secondary type 2 diabetes in patients with hypercortisolism. To do this, we ran simulations of OGTTs and recorded glucose and insulin responses under hypercortisolism while following the previously identified sequence of parameter values *σ*, v, and *I*_*m*_ describing the onset and progression of diabetes. The results are also shown in Fig. 6 (red curves), where we can see that excess CORT induces a shift of Starling’s curve toward higher fasting glucose levels, while preserving the same range of insulin responses to OGTTs.

## Discussion

Accounting for the dynamic interactions between the stress and metabolic axes is key to fully understand multisystemic conditions and to make sense of experimental and clinical observations subject to diurnal variability (*62, 66-69*). To advance this, we have developed a first-generation mathematical model that considers the dynamic interactions between glucocorticoid hormones and the mechanisms underpinning glucose metabolism. Our results suggest how these hormones modulate glucose and insulin responses to OGTTs with circadian and ultradian variability, and predict how hypercortisolism disrupts such variability. Moreover, our model provides plausible explanations regarding how CORT excess disrupts the mechanisms regulating glucose and insulin dynamics, and predicts what combination of mechanisms would be disrupted in the onset and progression of type 2 diabetes.

### Implications for diagnosis and treatment

OGTTs are a prevalent tool for evaluating glycaemic control and support the diagnosis and monitoring of metabolic disorders. Although current methods indicate measuring glucose levels at baseline and 2 hours post OGTT, there is growing recognition of the importance of considering additional time points, particularly for the early diagnosis of pre-diabetes (*70*). In line with this, our findings prompt for measuring the entire transient dynamics, with a higher sampling rate at least during the first hour to reliably locate the post-OGTT peak. As we have shown, mathematical methods can extract insights from rapid transient responses that would otherwise remain hidden if these data were not available. This is evident when examining the circadian and ultradian variability of glucose and insulin dynamic responses to OGTTs, which has significant importance in interpreting the results of this test in the context of the timing at which it was performed.

Our results also suggest OGTTs should be interpreted carefully in patients with diabetes secondary to glucocorticoid therapy or with endogenous cortisol disorders, with special attention to the transient post-OGTT dynamics. This is important in view of the fact that an OGTT may not be performed when a patient’s fasting glucose is normal, thus leading to an underestimation of the prevalence of diabetes in hypercortisolism (*15*). Our model is also a step towards an integrated theoretical framework that accounts for the circadian dynamics of insulin and glucose utilisation. Such approach is key to design novel clinical interventions, repurpose drugs, and implement chronotherapies for the treatment of diabetes secondary to hypercortisolism (*14, 71, 72*).

### Model reaches, limitations and future directions

As any mathematical model, ours attempts to strike a balance between accounting for enough physiological details for it to be meaningful, and simplifying the abstraction of such details to preserve its tractability. Whilst glucocorticoids have ubiquitous effects on the organism, for simplicity, we consider only the direct effects on glucose regulation. This allowed us to focus on the questions of how complex glucocorticoid rhythms modulate glucose and insulin dynamics, the effects of hypercortisolism on circadian variability, and the disruption of such mechanisms in the development of type 2 diabetes. In exchange, we omitted other glucocorticoid effects that may also contribute indirectly to metabolic disruptions. Obesity is perhaps the best example of this, as studies in mice demonstrate that abnormal glucocorticoid dynamics can increase the accumulation of fat mass (*67*). Whether such a hormone signalling filtering mechanism occurs in humans and its implications in regulating energy metabolism remains to be explored. Similarly, mounting evidence suggests that circadian clock genes within the brain, skeletal muscle, adipose tissue, gut, liver and pancreas (to name a few) may play a major role in regulating insulin sensitivity, possibly cooperating with glucocorticoids in controlling circadian variability (*2, 73, 74*). We envisage that the next generation of mathematical models will need to account for at least some of these long-term dynamic changes encoded at the gene and tissue level to develop a systemic understanding of chronodisruption.

Amongst the several areas of opportunity to improve the model, considering the dynamics of glucose and glucocorticoids across different tissues is perhaps the most promising because of its physiological importance. For example, in the model we assume that the bulk of glucose utilisation occurs in skeletal muscle and fat tissue, while leaving out the brain, the gut and other organs. Similarly, we focus the analysis on glucose, insulin and glucocorticoid levels in blood because of their clinical relevance, although this implicitly assumes blood levels are a proxy of the glucose dynamics within organs. We recognise that a better approach would be to use data on hormone and glucose levels collected directly from different tissues (e.g., muscle and fat), and modern sampling techniques are able to do this with high temporal resolution (*75, 76*). Considering the model limitations, our results show that subtle dynamic effects following OGTTs are important to properly characterise chronodisruption, and such effects can only be detected through high frequency sampling techniques.

Finally, to better understand the effects of glucocorticoid agonists and antagonists in the context of circadian glucose metabolism, we need to develop human studies that combine experimental and modelling analysis on both short and long term dynamic responses. Likewise, designing chronotherapies for stress-related metabolic conditions will demand a theoretical framework that integrates glucocorticoid rhythms, their effects on glucose dynamics and vice versa (*77*). Our model is a step toward such an integrative theoretical framework, and constitutes a shift from a descriptive to a mechanistic understanding of the dynamic links between stress and metabolism.

## General

The authors thank Prof John Terry for travel support via the UKRI Global Challenges Research Fund.

## Author contributions

E.Z., K.C.A.W., K.T.A. and M.A.H.V. conceived and designed the study. E.Z., C.A.G.G., K.C.A.W., K.T.A. and M.A.H.V. developed the mathematical model. C.A.G.G., E.Z., R.B. and M.A.H.V. performed the analytical calculations and computer simulations. E.Z., K.C.A.W., K.T.A. and M.A.H.V. supervised the project. E.Z. wrote the manuscript with support of C.A.G.G., K.C.A.W., K.T.A. and M.A.H.V.

