## Supplementary Material for "Dynamic modulation of glucose utilisation by glucocorticoid rhythms in health and disease"

### 1 Model Development

Glucose uptake by cells depends on a familiy of glucose transport membrane proteins, GLUTs, with GLUT1-4 isoforms posessing well-known roles as glucose transporters in different cell types [Thorens and Mueckler, 2009; Mueckler and Thorens, 2013]. The total glucose uptake per unit of time can be estimated as the sum of the average whole-body glucose uptake mediated by GLUT1-4. However, different cell types present different distributions of GLUTs in their cell membranes, with each GLUT possessing a different affinity for extracellular glucose. This is often represented via their associated Michaelis-Menten constant $K_m$ [Joost and Thorens, 2001; Unger, 1991]. The differential affinity poises cells to respond selectively to different plasma glucose concentrations, from normal physiological dynamic ranges (e.g., fasting and feeding) to pathological (e.g., diabetes and other metabolic disorders). In general, GLUT1 and GLUT3 transporters can be regarded as high affinity transporters (low $K_m$) that provide basic glucose supply to most cells, whereas GLUT4 contribute differentially to glucose uptake in muscle cells and adypocytes through a process dependent on insulin, glucocorticoids, and other factors. Conversely, GLUT2 transporters have a lower affinity for glucose (high $K_m$) and are highly expressed in $\beta$ cells and liver, where they are regarded as glucose sensors that mediate insulin secretion by $\beta$ cells and trigger glycogenolysis in the liver [Unger, 1991; Scheepers et al., 2004].

In adypocytes and muscle cells, GLUT1, GLUT3 and GLUT4 amplify glucose uptake via an insulin-dependent mechanism that translocates these transporters from intracellular pools to the cell membrane [Scheepers et al., 2004; Wilson et al., 1995]. This process is antagonised by glucocorticoids such as dexamethasone, which translocates GLUTs from the plasma membrane back to intracellular compartments [Ngo et al., 2009; Palmada et al., 2006], whereas insulin lowers the effect of dexamethasone at high concentrations [Carter-Su and Okamoto, 1987; Horner et al., 1987]. As suggested by the experiments performed by Carter-Su and Okamoto [1987] on adipocytes, the concentration of glucose transporters in the membrane is strongly correlated with the rate of glucose transport. In particular, the profiles of glucose transport and the labeled membrane transporter as functions of insulin concentration and dexamethasone are similar.

#### 1.1 Model equations

Let $G$ and $I$ represent the circulating concentrations of glucose and insulin, respectively. The change in $G$ over time can be thought of as the sum of glucose boluses from external sources (e.g., meals and OGTTs), endogenous glucose production via gluconeogenesis and glycogenolysis, minus glucose uptake by cells and a degradation term (Fig. 1). The change in $I$ over time is largely governed by pancreatic insulin secretion, which in turn depends on glucose sensing and is antagonised by glucocorticoids, minus a degradation term. To model the antagonistic effects of insulin and glucocorticoids, we assume $T \in (0,1)$ represents the fraction of translocatable GLUTs in the plasma membrane of peripheral cells, and $1-T$ the fraction of intracellularly docked transporters. Dynamic changes in insulin and glucocorticoids thus redistribute these fractions by translocating GLUTs between intracellular compartments and the plasma membrane. The model equations are as follows:

$$\dot{G}(t) = G_i - aU_c(G)T - r_G G, \quad (1)$$
$$\dot{I}(t) = \epsilon + \sigma S_\beta(G) h_Q(Q) - r_I I, \quad (2)$$
$$\dot{T}(t) = \big(u + v_I f_I(I)\big)(1-T) - \big(d + v_Q f_Q(Q)\big)T. \quad (3)$$

where the glucose input $G_i$ sums the contribution of glucose boluses $F(t)$ due to feeding or OGTTs and the endogenous glucose production $v f_e(G)$ due to gluconeogenesis and glycogenolysis, with $v$ as the maximum

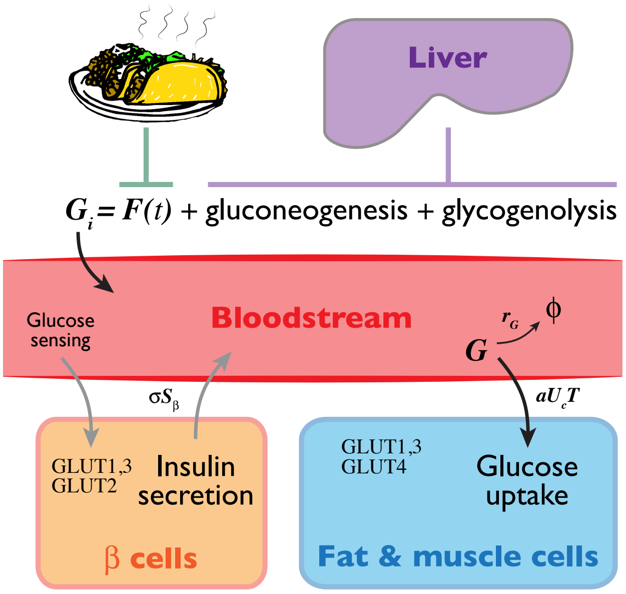


Figure 1: ***A systems level view of glucose utilisation.*** *Blood glucose concentration is maintained at basal levels by gluconeogenesis and glycogenolysis during fasting, increases transiently due to feeding (or OGTTs), and decreases due to cellular uptake and degradation. Circulating levels of glucose are also sensed by pancreatic $\beta$ cells, which in normal conditions secrete insulin in response to high levels of glucose (e.g., typically after feeding), a process that amplifies glucose uptake by peripheral cells.*

rate for such process:

$$G_i \quad = \quad F(t) + vf_e(G) \tag{4}$$

$aU_c(G)T$ accounts for the glucose uptake by fat and muscle cells, depending on the fraction of active GLUTs (denoted by $T$) and limited by the maximum uptake rate $a$. These tissues possess glucose transport systems with low and medium half-maximum constants, so the total glucose uptake can be represented as a weighted sum of the glucose transport mediated by a combination of GLUTs with low and medium activation thresholds. Thus,

$$U_c(G) \quad = \quad \underbrace{c_L f_L(G)}_{\text{GLUT1,3}} + \underbrace{c_M f_M(G)}_{\text{GLUT4}} \tag{5}$$

where $c_L$ and $c_M$ account for the relative contribution to glucose uptake by low and medium $K_m$ GLUTs in cell membranes. $r_G G$ accounts for first order glucose removal.

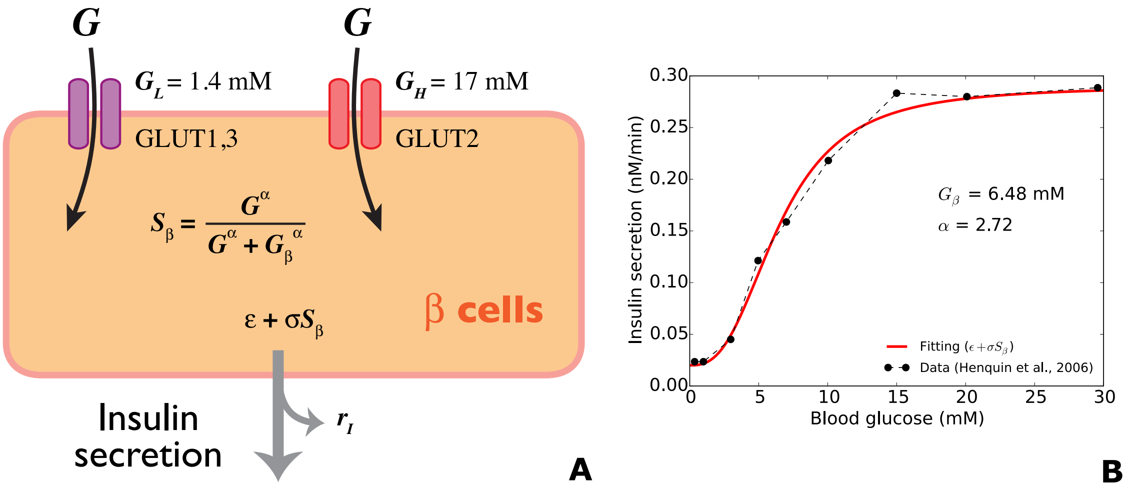


Figure 2: ***Glucose sensing and insulin secretion in $\beta$ cells. A****. Glucose sensing in $\beta$ cells is mediated by GLUT1, GLUT2 and GLUT3 transporters.* ***B****. Sensed levels of glucose trigger insulin secretion in a non-linear way, which is modelled via a sigmoidal Hill-type function $S_\beta$ fitted to data [Henquin et al., 2006].*

$S_\beta(G)$ is an expression accounting for glucose sensing in $\beta$ cells, which is mediated by GLUT1, GLUT2 and GLUT3, and was modelled as a sigmoidally increasing function of glucose and fitted to data [Henquin et al., 2006] (Fig. 2). This included a basal insulin secretion rate $\epsilon$ and a maximum insulin secretory rate $\sigma$. The factor $h_Q(Q)$ accounts for the modulatory effects of glucocorticoids on $\beta$ cell insulin secretion. $r_I I$ accounts for first order insulin removal.

Lastly, the equation representing the change in the fraction of active GLUTs over time (Eq. 3) can be better understood as composed by two terms on the right hand side, each representing a direction for GLUT translocation (Fig. 3). The first term accounts for those mechanisms that translocate GLUTs to the cell membrane $u + v_I f_I(I)$, and is thus factored by the fraction of GLUTs in intracellular pools $(1 - T)$. The second term accounts for those mechanisms that translocate GLUTs back to intracellular pools $d + v_Q f_Q(Q)$, and is thus factored by the fraction of GLUTs in the cell membrane $T$, which are the ones that actively transport glucose into cells. The mechanisms described within each term are further divided in two parts. In the first term, $u$ represents the basal translocation rate from intracellular pools up to the cell membrane, while $f_I(I)$ accounts for the insulin-dependent translocation in the same direction, at maximum rate $v_I$. In the second term, $d$ represents the basal translocation rate from the cell membrane down to intracellular pools, while $f_Q(Q)$ accounts for the glucocorticoid-dependent translocation in the same direction, at maximum rate $v_Q$.

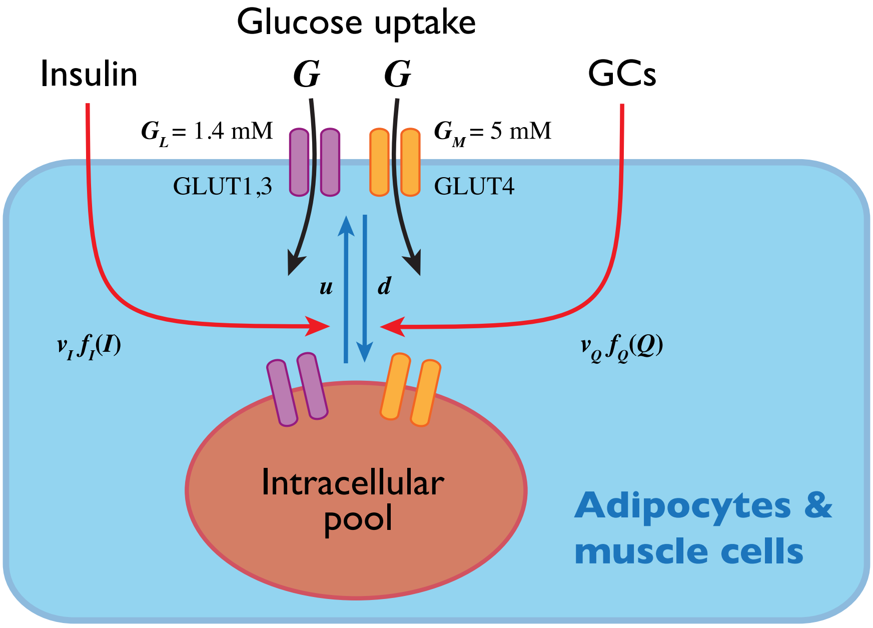


Figure 3: ***Glucose uptake and translocation of GLUTs in fat and muscle cells.*** *Glucose uptake into peripheral cells is mediated by GLUT1,3 and GLUT4 transporters. These transporters are distributed between the cell membrane and intracellular pools. While insulin promotes the translocation of GLUTs from intracellular pools to the cell membrane, glucocorticoids promote their translocation in the opposite direction. This creates a dynamic antagonism between insulin and glucocorticoids in regulating glucose uptake.*

The model in Eqs. 1-3 is supported by sigmoidal functions. These have the general form $\phi(x, K_m, h) = \dfrac{x^h}{x^h + K_m^h}$, where $K_m$ is the half maximum constant and $h$ is the Hill coefficient $(h > 1)$, with the special case of a Michaelis-Menten type mechanism when $h = 1$. In our model, these functions are:

$$f_e(G) = 1 - \phi(G, G_e, 1) \quad (6)$$
$$f_L(G) = \phi(G, G_L, 1) \quad (7)$$
$$f_M(G) = \phi(G, G_M, 1) \quad (8)$$
$$S_\beta(G) = \phi(G, G_\beta, \alpha) \quad (9)$$
$$f_I(I) = \phi(I, I_M, 2) \quad (10)$$
$$f_Q(Q) = \phi(\omega_f Q, Q_M, 2) + D_f \quad (11)$$
$$h_Q(Q) = 1 - \phi(\omega_h Q, k_Q, n) + D_h \quad (12)$$

In Eq. 6, we have used a decreasing Hill type function to account for the increase in gluconeogenesis and glycogenolysis following glucose depletion, which is the scenario observed during fasting [Unger, 1991]. The functions in Eqs. 7-8 account for the Michaelis-Menten glucose uptake into cells at low and medium $K_m$ (Fig. 3), whereas Eq. 9 accounts for glucose sensing in $\beta$ cells leading to insulin secretion (Fig. 2B). Lastly, the functions in Eqs. 10-11 account for insulin and glucocorticoid dependent translocation of GLUTs. Both Eqs. 11 and 12 have additional control parameters ($\omega_f$, $D_f$ and $\omega_h$, $D_h$, respectively) that account for the effects of dexamethasone shutting down the HPA axis ($\omega_f, \omega_h \ll 1$) with simultaneous sustained activation of the glucocorticoid receptor ($D_f, D_h > 0$). Note that Eq. 12 is a decreasing function of glucocorticoids, which accounts for its inhibitory effects on pancreatic insulin secretion.

### 1.2 Model drives

In addition to the governing Eqs. 1-3, we must also consider dynamic variables driving the system time evolution. For instance, fasting glucose levels in blood range between 80 to 100 mg/dl (equivalent to roughly 4.5 to 5.5 mM)[1] but increase sharply after meals. In healthy individuals, this causes insulin levels to rise (peak occurs ∼30 min after an oral glucose tolerance test (OGTT) [Kautzky-Willer et al., 1996]), which in turn increases the fraction of active GLUT transporters in fat and muscle cells. Regarding glucocorticoids, plasma levels of cortisol in healthy humans fluctuate within a range of 0.5 - 5 $\mu$M. These fluctuations exhibit ultradian periodicity ($T_u \approx 75$ min) with a circadianly-modulated amplitude that peaks early in the morning and decreases during the day, before rising again during the night [Spiga et al., 2015]. One of the purposes of the model is to explore how these endogenous oscillations modulate insulin sensitivity and glucose uptake in peripheral tissues, mediated by dynamically regulating the fraction of GLUT transporters in cell membranes.

To account for glucose boluses originated from OGTTs we used the pulse-like function $F(t)$:

$$F(t) = A\frac{t}{\tau}e^{(1-\frac{t}{\tau})} \quad (13)$$

where $A$ is the peak amplitude of $F(t)$ and $\tau$ is the time at which such a peak is reached. These parameters were fitted to match the timescale and peak value of blood glucose surges from baseline levels (Fig. 4A) [Kautzky-Willer et al., 1996].

Similarly, the external drive by glucocorticoids can be accounted for by a function $Q(t)$ representing ultradian oscillations with circadianly modulated amplitude:

$$Q(t) = A_G sin^2\left(\frac{\pi t}{T_c}\right) sin^2\left(\frac{\pi t}{T_u}\right) + A_m sin^2\left(\frac{\pi t}{T_c}\right) + B \quad (14)$$

Due to the interindividual variability of glucocorticoid dynamics, we manually fitted the parameters in Eq. 14 to reproduce the average amplitudes and periodicity observed in humans (Fig. 4B) [Spiga et al., 2015].

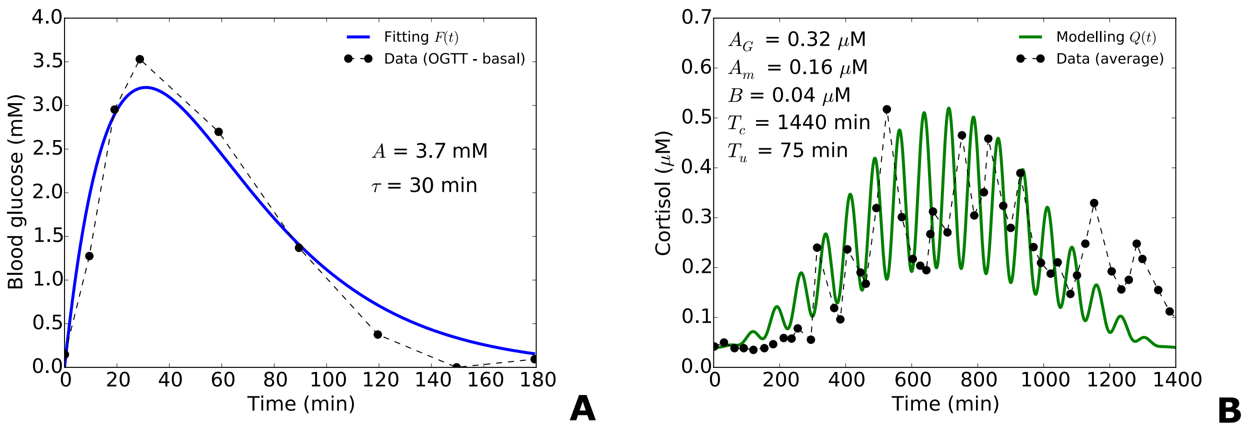


Figure 4: ***Model drives. A****. The function $F(t)$ models the pulse-like behaviour resulting from a glucose bolus. We fitted its parameters to reproduce a typical blood glucose surge resulting from an OGTT [Kautzky-Willer et al., 1996].* ***B****. The function $Q(t)$ models average cortisol fluctuations in humans [Spiga et al., 2015]. Time zero corresponds to 7 p.m. when cortisol levels are at its minimum, then increase progressively during the night until reaching a maximum at 7 a.m. and decreases again during the day.*

Lastly, we simulated idealised scenarios of hypercortisolism and the effects of an HPA agonist as illustrated in Fig. 5.

[1] 1 mM glucose is equivalent to a mass concentration of 18 mg/dl.

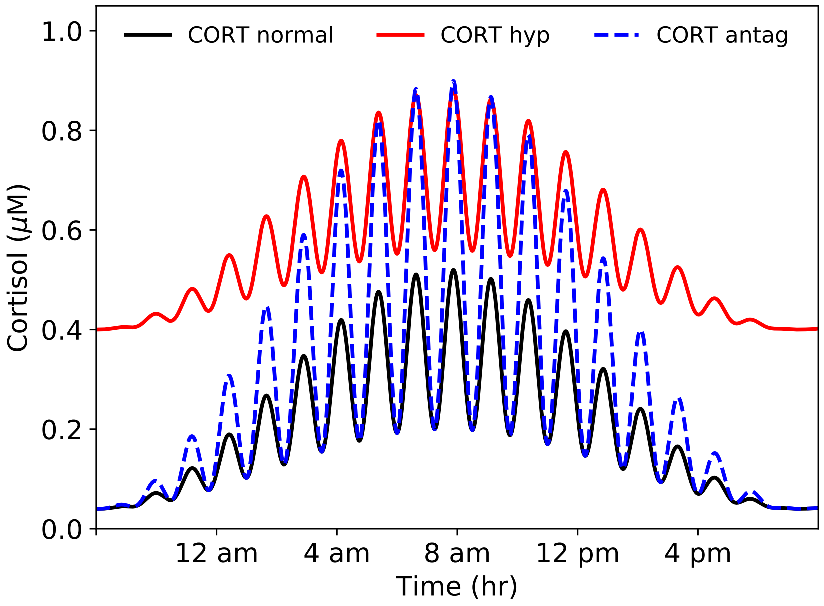


Figure 5: ***Cortisol dynamics.*** *Ultradian oscillations of cortisol with circadianly modulated amplitude simulated for the normal physiological scenario (black), hypercortisolism (red), and under the hypothetical effect of an antagonist (blue).*

### 2 Parameter Estimation and Fitting

The parameters described in this section are summarised in Table 2. We prioritised using values for humans directly reported in the literature, followed by those estimated from experiments in humans or animal cell lines. Where this was not possible we fixed values arbitrarily within biologically plausible ranges and fitted to data.

**Estimation of $u$, $d$, $v_I$ and $v_Q$.** To estimate the kinetic rates associated to GLUT translocation we used data from experiments performed on a rat skeletal muscle cell line that determined the basal and insulin-dependent distribution of GLUT1, GLUT3 and GLUT4 [Wilson et al., 1995]. In these experiments, the authors estimated the cell membrane levels of each GLUT protein following 30 min of insulin treatment (100 nM) vs non-treated controls. We have summarised these results in Table 1, showing concentrations for each cellular compartment analysed (membrane and intracellular microsomes) under two different conditions (control vs insulin stimulated).

| **Transporter** | **Membrane** | | **Microsomes** | |
|---|---|---|---|---|
| | **Control** | **Insulin** | **Control** | **Insulin** |
| GLUT1 | 0.014 | 0.017 | 0.0053 | 0.0029 |
| GLUT3 | 0.0097 | 0.014 | 0.0046 | 0.0018 |
| GLUT4 | 0.012 | 0.015 | 0.0061 | 0.0044 |
| **Total** | **0.0357** | **0.046** | **0.016** | **0.0091** |

Table 1: *Concentration [pmol/mg] of GLUT proteins in the cell membrane and intracellular microsomes, under insulin stimulation vs controls, estimated from [Wilson et al., 1995]*

From Table 1, we can calculate the total concentration of GLUT1, GLUT3 and GLUT4 in the cell by adding the concentration of each transporter in the membrane and microsomes for both experimental conditions and calculating the mean value, resulting in $[GLUT]_{total} = 0.0534$ pmol/mg. We also note that the incubation time in the experiments made by [Wilson et al., 1995] is long enough to consider their results as steady state distributions of GLUTs. This means we can re-write Eq. 3 as:

$$\dot{T}(t) \quad = \quad \frac{T_\infty(I,Q) - T}{\tau(I,Q)} \qquad (15)$$

where $T_\infty(I,Q)$ and $\tau(I,Q)$ can be interpreted as the steady state and the time constant for $T$, respectively.

These are given by:

$$T_\infty(I,Q) = \frac{u + v_I f_I(I)}{u + v_I f_I(I) + d + v_Q f_Q(Q)}, \quad (16)$$

$$\tau(I,Q) = \frac{1}{u + v_I f_I(I) + d + v_Q f_Q(Q)}. \quad (17)$$

Written in this way, the proportion $T$ of GLUT transporters in the membrane will eventually reach the steady state value $T_\infty(I,Q)$ at a rate determined by the time constant $\tau(I,Q)$. If we only consider insulin effects (i.e., $Q \to 0$), the system will converge toward a higher steady state $T_\infty(I,0)$, whereas adding only glucocorticoids (i.e., $I \to 0$) will cause $T_\infty(0,Q)$ to shift to lower values. The time scale $\tau(I,Q)$ of these processes would thus depend on the balance between $v_I$ and $v_Q$.

If we assume the concentrations of insulin and glucocorticoids are at its maximum or minimum in a given incubation experiment, then functions in Eqs. 10 and 11 will converge to approximately 1 or 0, respectively. This leads to four possible steady states:

$$T_\infty(0,0) = \frac{u}{u+d}, \quad (18)$$

$$T_\infty(I_{max},0) = \frac{u+v_I}{u+v_I+d}, \quad (19)$$

$$T_\infty(0,Q_{max}) = \frac{u}{u+d+v_Q}, \quad (20)$$

$$T_\infty(I_{max},Q_{max}) = \frac{u+v_I}{u+v_I+d+v_Q}. \quad (21)$$

The values of the first two steady states can be estimated directly from the distributions reported in Table 1:

$$T_\infty(0,0) = \frac{[GLUT]^{control}_{mem}}{[GLUT]_{total}} = \frac{0.0357\ pmol/mg}{0.0534\ pmol/mg} = 0.67 \quad (22)$$

$$T_\infty(I_{max},0) = \frac{[GLUT]^{insulin}_{mem}}{[GLUT]_{total}} = \frac{0.046\ pmol/mg}{0.0534\ pmol/mg} = 0.87 \quad (23)$$

From Eqs. 18 and 22, we can see that $u \approx 2d$. Moreover, the rate of GLUT4 exocytosis in human muscle cells in basal conditions ($I = 0$ and $Q = 0$) has been measured (Fig. 5C of [Karlsson et al., 2009]), which we take as the value for $u = 0.01$ min$^{-1}$. It immediately follows that $d = 0.0049$ min$^{-1}$.

Experiments by [Carter-Su and Okamoto, 1987] in rat adipocytes show convergence of glucose dependent uptake to a steady state following insulin stimulation, with or without previous incubation with dexamethasone. Assuming these results reflect the abundance of GLUTs in the membrane, and keeping in mind the glucose levels explored ranged between 0 - 10 mM, it is possible to estimate the GLUT fraction in the membrane when cells were simultaneously incubated with insulin and dexamethasone ($T_\infty(I_{max},Q_{max})$), and compare it when only insulin was used ($T_\infty(I_{max},0)$). Following this, a graphical inspection of Fig. 3 in [Carter-Su and Okamoto, 1987] allows us to estimate $T_\infty(I_{max},Q_{max}) = 0.64$.

Then, from Eq. 19:

$$v_I = \frac{u - (u+d)T_\infty(I_{max},0)}{T_\infty(I_{max},0) - 1}, \quad (24)$$

and from Eq. 21:

$$v_Q = \frac{u+v_I}{T_\infty(I_{max},Q_{max})} - (u+v_I+d). \quad (25)$$

Since all the parameters in the right hand side of Eqs. 24 and 25 are now estimated, we can calculate $v_I$ = 0.023 min$^{-1}$ and $v_Q$ = 0.014 min$^{-1}$.

**Estimation of $c_L$ and $c_M$.** To estimate these relative contributions, we just need to keep in mind that $c_L$ + $c_M$ = 1, and look at the concentrations reported by [Wilson et al., 1995] for low $K_m$ transporters (GLUT1 and GLUT3) and high $K_m$ transporters (GLUT4) in the cell membrane of rat muscle cells (summarised in Table 1). From this, we can easily estimate $c_L$ = 0.66 and $c_M$ = 0.34.

| Parameter | Value | Description | Source |
|---|---|---|---|
| $v$ | 18.69 mM·min$^{-1}$ | Maximum glucose endogenous production due to gluconeogenesis and glycogenolysis. | [Kautzky-Willer et al., 1996][(3)] |
| $a$ | 17.97 mM · min$^{-1}$ | Glucose absorption rate in cells. | [Kautzky-Willer et al., 1996][(3)] |
| $u$ | 0.01 min$^{-1}$ | Insulin and glucocorticoid independent translocation rate of GLUTs from intracellular pools to the plasma membrane. | [Karlsson et al., 2009][(1)] |
| $d$ | 0.0049 min$^{-1}$ | Insulin and glucocorticoid independent translocation rate of GLUTs from the plasma membrane to intracellular pools. | [Wilson et al., 1995][(2)] |
| $v_I$ | 0.023 min$^{-1}$ | Insulin dependent maximum translocation rate of GLUTs from intracellular pools to the plasma membrane. | [Wilson et al., 1995][(2)] |
| $v_Q$ | 0.014 min$^{-1}$ | Glucocorticoid dependent maximum translocation rate of GLUTs from the plasma membrane to intracellular pools. | [Wilson et al., 1995][(2)] |
| $c_L$ | 0.66 | Proportion of low $K_m$ GLUT transporters in peripheral cells. | [Wilson et al., 1995][(2)] |
| $c_M$ | 0.34 | Proportion of medium $K_m$ GLUT transporters in peripheral cells. | [Wilson et al., 1995][(2)] |
| $\epsilon$ | 0.02 nM · min $^{-1}$ | Basal insulin secretion rate during fasting. | [Henquin et al., 2006][(2)] |
| $\sigma$ | 0.29 nM · min$^{-1}$ | Glucose dependent maximum insulin secretion rate. | [Henquin et al., 2006] [Kautzky-Willer et al., 1996][(2)] |
| $r_I$ | 0.47 min$^{-1}$ | Insulin removal rate. | Arbitrary |
| $r_G$ | 0.2 min$^{-1}$ | Glucose removal rate. | Arbitrary |
| $G_e$ | 5 mM | Endogenous glucose production half maximum constant. | Arbitrary |
| $G_L$ | 1.4 mM | Low $K_M$ GLUT mediated glucose uptake. | [Unger, 1991][(1)] |
| $G_M$ | 4 mM | Medium $K_M$ GLUT mediated glucose uptake. | [Unger, 1991][(1)] |
| $I_M$ | 0.081 nM | Half maximum constant for insulin mediated translocation of GLUTs. | [Carter-Su and Okamoto, 1987][(2)] |
| $Q_M$ | 0.28 $\mu$M | Half maximum constant for glucocorticoid mediated translocation of GLUTs. | [Carter-Su and Okamoto, 1987][(2)] |
| $G_\beta$ | 6.48 mM | Half maximum constant of glucose dependent insulin secretion. | [Henquin et al., 2006][(3)] |
| $\alpha$ | 2.72 | Hill coefficient of glucose dependent insulin secretion. | [Henquin et al., 2006][(3)] |

Table 2: *Model parameter values. (1) Reported. (2) Estimated. (3) Fitted to data.*

**Estimation of $I_M$.** This parameter was estimated from experiments that measured glucose uptake following insulin incubation of rat adipocytes [Carter-Su and Okamoto, 1987]. The range of insulin stimulation was 0 - 2000 $\mu$U/ml, increasing the uptake of the non-metabolizable glucose analogue 3-OMG from 3.1 - 40 $\mu$mol/seg · L. From the data described in Fig. 2 of [Carter-Su and Okamoto, 1987], it was possible to interpolate at which concentration insulin exerts half its maximum effect, which we express in Molar units as $I_M = 0.081$ nM.

**Estimation of $Q_M$.** It is mentioned in [Carter-Su and Okamoto, 1987] that a concentration of 0.1 $\mu$M of dexamethasone maximally inhibits GLUT translocation to the membrane of rat adipocytes. Following this, we have approximated the half maximum constant to $Q_M = 0.05$ $\mu$M.

**Estimation of $\sigma$ and $\epsilon$.** Acording to [Henquin et al., 2017], the pancreas of a lean non-diabetic person contains ∼10.29 mg of insulin, with only a small fraction secreted in response to glucose. In fact, [Henquin et al., 2006] suggest that the pancreas of a healthy adult releases ∼0.078% of its insulin content per minute at a maximal glucose stimulation of 30 mM. This translates to a maximal secretion rate of 8.03 $\mu$g/min, which we can express as 1.38 nM/min using the molecular weight of insulin (5808 Da). Now, since we're interested in the maximum production rate in plasma, we should consider the blood volume in which it is diluted. To do this, we considered $\sim$4.82 L of blood in humans, which then gives a maximal insulin production of $\sigma$ = 0.29 nM/min.

On the other hand, we can see from Fig. 1B in [Henquin et al., 2006] that at near zero glucose stimulation, there is continued basal insulin secretion amounting to $\sim$8% of maximum insulin production. This allows us to estimate $\epsilon = 0.02$ nM/min.

**Remaining parameters.** The remaining parameter values have been either reported directly in the literature ($G_L = 1.4$ mM, $G_M = 5$ mM) [Unger, 1991], fixed arbitrarily ($r_G = 0.01$ min$^{-1}$, $r_I = 0.5$ min$^{-1}$, $G_e$ = 4 mM), or fitted to published data ($G_\beta = 6.48$ mM, $\alpha = 2.72$, $v = 15.12$ mM/min, $a = 16.27$ mM/min) [Henquin et al., 2006; Kautzky-Willer et al., 1996].
